# Rhes, a Striatal Enriched Protein, Regulates Post-Translational Small-Ubiquitin-like-Modifier (SUMO) Modification of Nuclear Proteins and Alters Gene Expression

**DOI:** 10.1101/2020.06.18.160044

**Authors:** Oscar Rivera, Manish Sharma, Neelam Shahani, Uri Nimrod Ramírez-Jarquín, Gogce Crynen, Pabalu Karunadharma, Francis McManus, Thibault Pierre, Srinivasa Subramaniam

**Affiliations:** Department of Neuroscience, The Scripps Research Institute, Florida, USA, 33458; Center for Computational Biology and Bioinformatics, The Scripps Research Institute Jupiter, FL. 33458, USA; The Scripps Research Institute, Genomic Core, Jupiter, FL 33458, USA; Institute for Research in Immunology and Cancer, Université de Montréal, Montréal, H3T 1J4, Canada; Department of Chemistry, Université de Montréal, Montréal, Québec, Canada

**Keywords:** Medium spiny neurons, SUMO E3-ligase, Gene regulation, Brain, Differentiation, Morphology, Signaling, Perinuclear membrane

## Abstract

Rhes (Ras homolog enriched in the striatum) is a multifunctional protein that orchestrates striatal toxicity, motor behaviors and abnormal movements associated with dopaminergic signaling, Huntington disease and Parkinson disease signaling in the striatum. Rhes engineers membranous tunneling nanotube-like structures and promotes intercellular protein and cargoes transport. Recent study revealed Rhes also regulates mitophagy via the Nix receptor. Despite these studies, the mechanisms through which Rhes mediates these diverse functions remains unclear. Rhes belongs to a small GTPase family member and consists of a unique C-terminal Small Ubiquitin-like Modifier (SUMO) E3-like domain that promotes the post-translational modification (PTM) of proteins with SUMO (SUMOylation) by promoting “cross-SUMOylation” of SUMO enzymes SUMO E1 (Aos1/Uba2) and SUMO E2 ligase (Ubc-9). However, the identity of the SUMO substrates of Rhes remains largely unknown. By combining high throughput interactome and SUMO proteomics we report that Rhes regulates the SUMOylation of nuclear proteins that are involved in the regulation of gene transcription. While Rhes has increased the SUMOylation of histone deacetylase 1 (HDAC1) and histone 2B, it had decreased the SUMOylation of heterogeneous nuclear ribonucleoprotein M (HNRNPM), protein polybromo-1 (PBRM1) and E3 SUMO-protein ligase (PIASy). We also found that Rhes itself is SUMOylated at 5 different lysine residues (K32, K110, K114, K120, K124 and K245). Furthermore, we found that Rhes regulates the expression of genes involved in cellular morphogenesis and differentiation in the striatum, in a SUMO-dependent manner. Our findings thus provide a previously undescribed role for Rhes in regulating SUMOylation of nuclear targets and in orchestrating striatal gene expression via the SUMOylation.

## INTRODUCTION

Rhes (Ras homolog enriched in the striatum) mRNA is highly expressed in the dopamine D1 receptor (D1R)- or D2R-expressing medium spiny neurons (MSNs) and cholinergic interneurons in the striatum; it is also expressed to some extent in other brain regions such as cortex and hippocampus [1–3]. Rhes is induced by thyroid hormones and can inhibit the cAMP/PKA pathway and N-type Ca^2+^channels (Cav 2.2) [4–7]. We have found several new roles for Rhes in the striatum. Rhes can directly bind to, and activate, mTOR in a GTP-dependent manner, promoting L-DOPA-induced dyskinesia (LID) in a pre-clinical model of Parkinson’s disease [8]. Rhes also inhibits striatal motor activity through a ‘Rhesactome”—a protein network in the striatum—via its guanine nucleotide exchange factor (GEF), RasGRP1 [9], which we recently found also promotes LID [10]. Rhes affects autophagy via Beclin1, independent of mTOR signaling [11]. We also demonstrated a critical role for Rhes in Huntington disease (HD). We found Rhes interacts with mHTT, the genetic risk factor of HD, and makes it more soluble by virtue of its SUMOylation [9, 12]. Thus, Rhes and mHTT interaction increases cellular toxicity, a finding which is consistent with independent studies [12–19]. One report indicates that Rhes overexpression can be protective in HD [20], and so Rhes was tagged on its C-terminal end in this study to promote its mislocalization to the nucleus [12]. Rhes also plays a critical role in mutant tau-mediated pathology [21]. In addition, a rare, highly conserved de novo mutation (R57H) in Rhes was detected in twins diagnosed on the autistic spectrum [22].

Recently, tubular or tunneling nanotubes (TNTs), the fragile and inconspicuous membranous structures ranging from 50-200 nm diameter in size and 5-125 μm in length connecting two cells, have been reported in diverse cell types [23–39]. We have discovered that Rhes promotes TNT-like membranous protrusion called “Rhes tunnel” in multiple cell types, including primary striatal neurons [40]. Using mutagenesis, we found that the membrane binding site of Rhes (C263) is critical for Rhes tunnels. Mutagenesis also demonstrated a distinct role for the GTPase domain (1-171aa) and SUMO E3 ligase domain of Rhes (171-266aa) [40]. While the GTPase domain is defective in producing TNT-like protrusions, the SUMO E3 ligase domain induces TNTs, but shows diminished potency of cell-to-cell transport [40]. This data indicates a distinct role for Rhes domain, while SUMO E3 ligase domain promote Rhes tunnel, the GTPase domain is necessary for entering into the acceptor cell via Rhes tunnel. Given that Rhes is the first brain-enriched protein shown to promote TNT-like protrusions, a further investigation into its mechanisms and roles would improve our understanding of TNT-like communications in brain.

Rhes is capable of executing more than one function, at molecular, cellular, and organismal levels. Multifunctional proteins like Rhes are not rare in biology. For example, through network topological information with protein annotations, it was estimated that around 430 of human proteins can be classed as extreme multifunctional proteins [41], including p53 and Cyt C [42, 43]. Despite its multiple functions and molecular complexity, fundamental knowledge about the mechanisms by which Rhes works or how those mechanisms selectively impair distinct striatal disorders remains partial. We reported that Rhes physiologically regulates SUMO modification in vivo and mechanistically Rhes promotes the “cross-SUMOylation” of E1 (Aos1) and E2 (Ubc9) enzymes [44]. However, the identity of SUMO substrates of Rhes remains largely unknown. Using biochemical, proteomics and RNA seq tools, we investigated the SUMO substrates of Rhes and its putative role in the striatal gene expression.

## RESULTS

### Rhes interacts with SUMOylated proteins and enhances SUMOylation in cultured cells

Like ubiquitin, SUMO (3 paralogues in vertebrates: SUMO1, SUMO2, and SUMO3) is a conserved ~10.5kDa protein modification that is covalently attached to lysine residues on multiple substrate proteins in a dynamic and reversible manner [45]. To identify SUMO substrates of Rhes, we used mSUMO3 construct in combination with a large scale unbiased proteomics approach for SUMOylation site identification. mSUMO3 has a mutation in the C-terminal that can be cleaved by trypsin resulting in a C-terminus penta-peptide, which is compatible for mass-spectrometry (MS) detection [46, 47]. We employed mSUMO3 in this study because its transient expression produced more abundant protein that may aid in SUMO substrate detection in HEK293 cells, compared to mSUMO1 or mSUMO2 (Fig. 1A). First, we hypothesized that the SUMO proteins that binds to Rhes are the potential SUMO targets of Rhes. To examine this hypothesis, we co-expressed GST-Rhes or GST (control) with His-tagged mSUMO3 in HEK293 cells and affinity purified Rhes using GST-glutathione-affinity column. We found GST-Rhes readily bound to proteins that are modified by mSUMO3 (Fig. 1B). The GST-Rhes affinity purified samples were subjected to MS analysis to identify and quantify the interacting partners. We found several proteins that are readily bound to Rhes, including RAC1, TSC2, VDAC1, PLEC, DDX3X and mTOR, previously identified interactors of Rhes (Fig. 1C, Data file S1)[8, 9].

**Fig 1.**
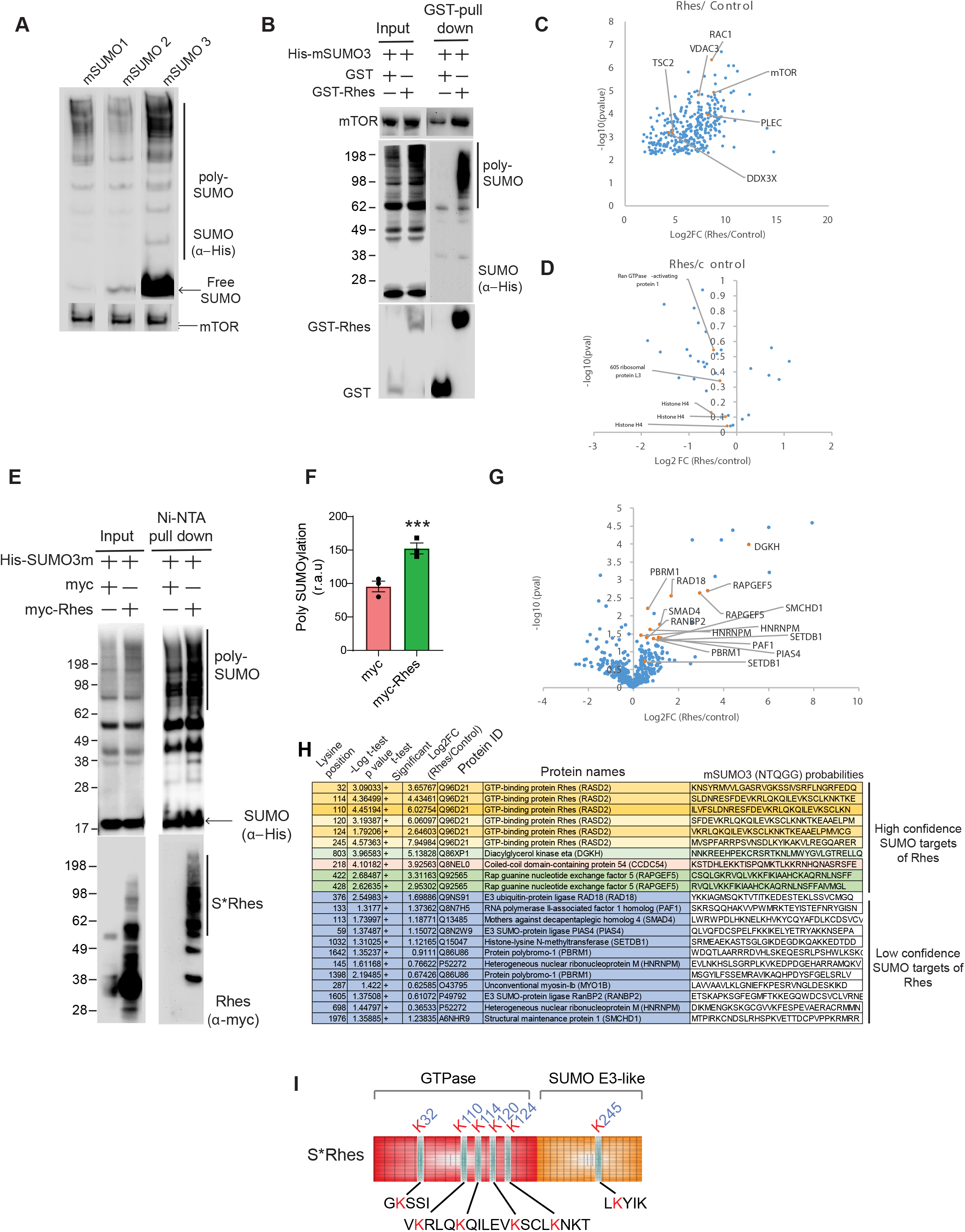
Interactome and SUMO proteome identifies putative SUMO substrates of Rhes. (**A**) Western blot of HEK293 cells expressing His-SUMO 1m, His-SUMO 2m, or His-SUMO 3m. (**B**) Western blot for indicated proteins after glutathione-affinity pulldown in HEK293 cells expressing GST + His-mSUMO3 (control) or GST-Rhes + His-mSUMO3 and corresponding input (5%) for the indicated proteins. (**C**) Volcano plot of proteins bound to affinity purified GST-Rhes (Rhes) co-expressing mSUMO3, compared to affinity purified GST co-expressing mSUMO3 (control), identified by LC-MS/MS in biological triplicate. (**D**) Volcano plot of SUMOylated proteins that are bound to glutathione enriched GST-Rhes (Rhes)+ His-mSUMO3, compared to glutathione enriched GST + His-mSUMO3 (control) vs. GST-Rhes co-expressing mSUMO3, identified by LC-MS/MS in biological triplicate. (**E**) Western blot of His-tagged SUMO enrichment using Ni-NTA TALON beads in HEK293 cells expressing myc + His-mSUMO3 or myc-Rhes + His-mSUMO3. S*Rhes represents SUMOylated Rhes. (**F**) Quantification of overall SUMOylation in Ni-NTA enriched, myc + His-mSUMO3 or myc-Rhes #x002B; His-mSUMO3. Data represents means ± SEM, (n =3), \*\*\**P <* 0.001, Student’s *t* test. (**G**) Volcano plot of SUMOylation site changes in myc-Rhes + His-mSUMO3 compared to myc + His-mSUMO3 (control), identified by LC-MS/MS in biological triplicate. (**H**) High and low confidence SUMO substrates of Rhes identified in *(G)*. (**I**) Representation of Rhes domains with mapping of the SUMO sites identified.

Attempts at identifying SUMOylated interacting partners from the GST pulldown were futile. Although we found targets such as RPL3 (K399), histone H4 (K6, K9, and K13) and RANGAP1 (K524) found SUMOylated, the log10 transformed p value for any of the targets did not reach below 0.05 (Fig. 1D) (Data file S2). We reasoned that the SUMOylated proteins that are bound to Rhes are either very few or below the threshold needed for the detection by MS. Moreover, GST pulldown experiments are conducted under native conditions, under which the SUMO specific proteases (SENPs) are highly active and could deconjugate the SUMO moiety from the target proteins[48]. Therefore, we resorted to a denaturing purification protocol to enrich SUMOylated proteins. We co-expressed myc-Rhes or myc (control) with His-tagged mSUMO3 in HEK293 cells and lysed the cells in guanidine-hydrochloride/urea denaturing buffer, followed by purification in Ni-NTA column. We found that the myc-Rhes significantly enhanced the overall SUMOylation of proteins (Fig. 1E, F) and Rhes was itself SUMOylated (S* Rhes, Fig. 1E), consistent with our previous report [12].

Together these results indicate a) Rhes interacts with multiple proteins, and b) Rhes is SUMOylated, and c) Rhes enhances the SUMOylation of several targets.

### Mass spectrometry reveals nuclear SUMO targets and SUMO modified lysines of Rhes

To identify the SUMO targets of Rhes, we subjected Ni-NTA enriched myc and myc-Rhes samples (Fig. 1E) to SUMO peptide immunopurifications using a custom antibody that recognizes the NQTGG remnant created on the modified lysine residue upon trypsin digestions. SUMOylation site identification and quantified were obtained by MS analysis. This analysis revealed numerous SUMOylated proteins and a significant increase of SUMOylation of at least 13 targets in myc-Rhes expressing cells compared to myc control (Fig. 1G, Data file S3).

Among the high-confidence SUMO substrates, we found Rhes increased the RapGEF5 SUMOylation at the lysine 422 and lysine 428 residues that are part of catalytic domain [49]. RapGEF5 is a direct target of 3’-5’-cyclic adenosine monophosphate (cAMP) and is involved in cAMP-mediated signal transduction through activation of the Ras-like small GTPase RAPs: RAP1A, RAP1B and RAP2A [50]. We found Rhes increased the SUMOylation at lysine 803 of diacylglycerol kinase eta (DGKH), an enzyme that generate phosphatidic acid (PA) and activates the Ras/B-Raf/C-Raf/MEK/ERK, We found Rhes also increased the SUMOylation of CCDC54 (the coiled coil domain containing protein 54), whose function is unknown, at lysine 318 (Fig. 1H).

Among the low confidence SUMO targets, we found that Rhes promoted the SUMOylation of the SUMO-3 ligases PIASy (K59) and RanBP2 (K1605), as well as protein polypbromo-1 (PBRM1, K1398, K1642), heterogenous nuclear ribonucleoprotein M (HNRNPM, K145, K698), histone-lysine N-methyltransferase (SETDB1, K1032), RNA polymerase II-associated factor 1 homolog (PAF1, K133) and mothers against decapentaplegic homolog 4 (SMAD4, K113) (Fig. 1H). Interestingly, Rhes is also one among the 4 high-confidence SUMOylated targets. Rhes was found SUMOylated at six lysine residues 32, 110, 114, 120, 124 and 245 that spanned across the protein. The GTPase domain of Rhes contains 5 SUMO modification sites and the C-terminal SUMO E3-like region harbors a SUMOylation site (Fig. 1I).

Together this data indicates that a) Rhes modulates SUMOylation of substrates involved in cellular signaling and nuclear functions and b) Rhes is SUMOylated at multiple lysine residues.

### Comparison of Rhes interactome and SUMOylation targets to further refine potential new SUMO targets of Rhes

Although the quantitative SUMO proteomic (Fig. 1) revealed novel putative SUMO targets of Rhes, the number of SUMO candidates identified are relatively low. Such results are not entirely unexpected because of the hurdles that are inherent to SUMOylation studies such as the low abundance of the SUMO modified targets and the highly dynamic nature of SUMOylation. Indeed, at any given time only <1% of the total protein are modified, further complicating their identification [51]. Finally, SUMOylation of certain substrates in the native tissues such as striatum may also be under the strict control of extracellular signaling.

Keeping all these limitations in perspective, we sought to identify the potential SUMO targets further, using an unbiased approach. So, we applied four different filtering to sort out SUMO substrates (Fig. 2A). First, we selected the common proteins by comparing Rhes interactors in HEK293 cells [Fig. 1C, (*a*)] with all the SUMOylated proteins in Rhes expressing HEK293 cells [Fig 1G, (*b*)] to obtain *a-1*. Second, we compared protein interactors of Rhes from the striatum [9] (*c*) with the SUMOylated proteins in Rhes expressing HEK293 cells (*b*) and obtained *b-1*. Third, we compared proteins interactor of Rhes in the striatum (*c*) with the SUMO interacting proteins in the brain [52] (*d*), and obtained *c-1.* Fourth, we sorted shared protein from *a-1*, *b-1* and *c-1* to obtain *d-1*. And fifth, we compared the high and low confidence SUMO substrates of Rhes (*e-1*) with proteins identified from (*d-1*) to obtain *f-1* (Data file S4). This approach has resulted in 27 novel targets that are SUMOylated as well as shown to interact with Rhes (Fig. 2B). Most of these targets, such as histone deacetylase 1 (HDAC1), protein inhibitor of activated STAT protein 4 (hPIASy or PIASy), histone-2B, (H2B), heterogeneous nuclear ribonucleoprotein M (HNRNPM) and polybromo 1 (PBRM1) have roles in gene regulation, indicating a strong nuclear function for Rhes.

**Fig 2.**
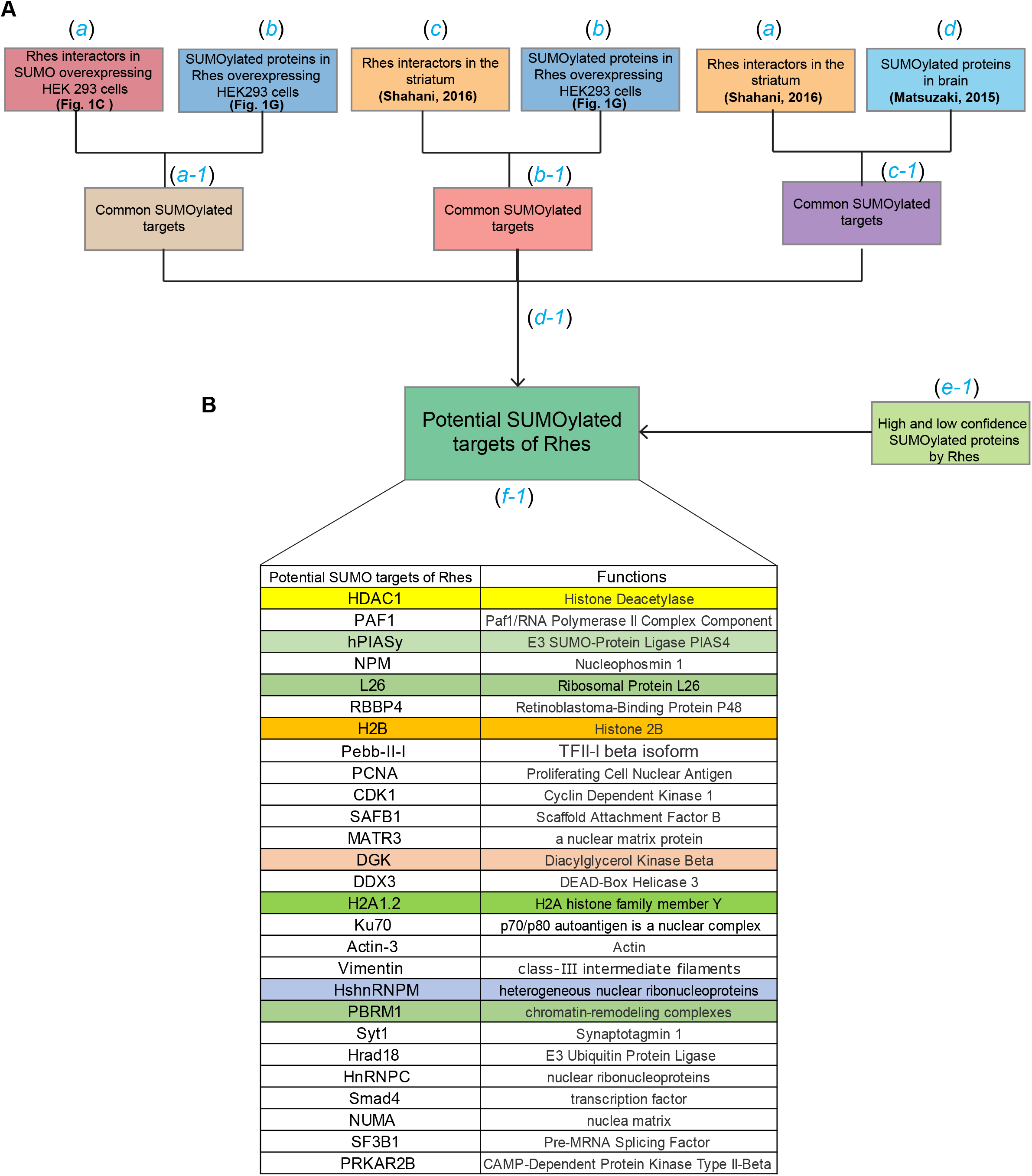
Flow chart to identify potential SUMOylated proteins. (**A**) Proteins that interacts with Rhes in His-mSUMO3 overexpression *(a)* are compared with all the SUMOylated proteins *(b)* and obtained *(a-1).* Rhes interactors in the striatum, Shahani, 2016[9] *(c)* are compared with *(b) to* obtain *(b-1).* Then, *(a)* is compared with SUMOylated proteins in the brain, Matsuzaki, 2015[52], *(d)* to obtain *(c-1).* Later, a-1, b-1 and c-1 are compared to obtain d-1. (**B**) Finally, the high and low confidence SUMO substrates of Rhes *(e-1)* were compared to d-1 to generate a list of the top 27 putative SUMO substrates of Rhes *(f-1).*

### Rhes regulates SUMOylation of nuclear targets in cells

Next we hypothesized that Rhes can directly modulate the SUMOylation of the targets identified in Fig 2. To test this hypothesis, we selected certain targets (color coded) based on the already available cDNA constructs from Addgene (Fig 2B). So, we carried out Ni-NTA denaturing protocol, (Fig. 3) for these nuclear targets by expressing them in in HEK293 cells.

**Fig 3.**
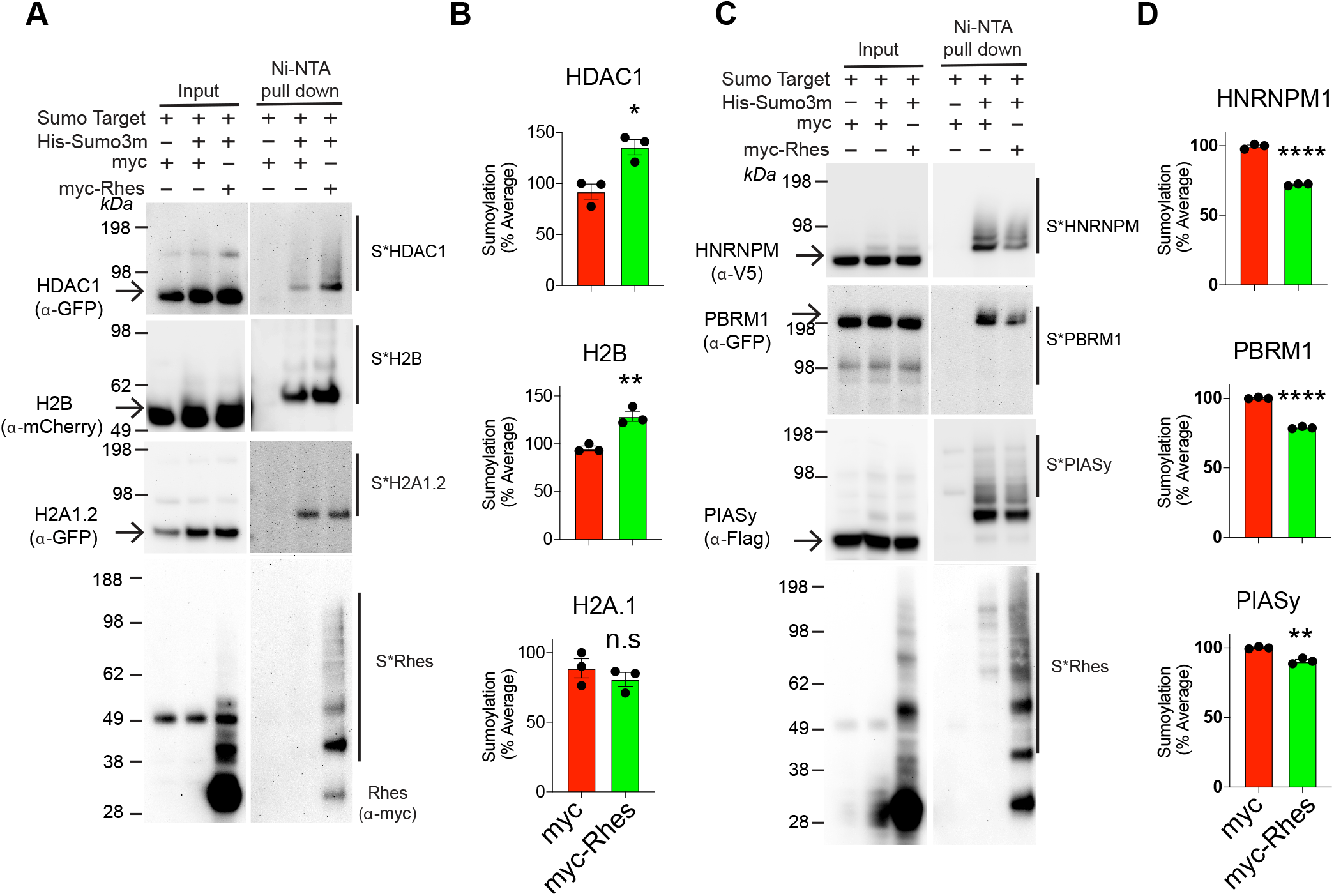
Rhes regulates the SUMOylation of nuclear targets. (**A**) Western blot of Ni-NTA enrichment of SUMOylated proteins (S*) from the lysates obtained from HEK293 cells expressing myc + His-mSUMO3 or myc-Rhes + His-mSUMO3 constructs either with GFP-HDAC1, m-cherry H2B or GFP-H2A1.2. (**B**). Bar graphs indicates quantification of SUMOylation (%) from (A). Data represents mean ± SEM, (n =3), \**P <* 0.05, \*\**P <* 0.01, Student’s *t* test. n.s: not significant. (**C**) Western blot of Ni-NTA enrichment of SUMOylated proteins from the lysates obtained from HEK293 cells expressing myc + His-mSUMO3 or myc-Rhes + His-mSUMO3 constructs either with V5-HNRNPM1, GFP-PBRM1 or Flag-PIASy. (**D**). Bar graphs indicates quantification of SUMOylation (%) from (C). Data represents mean ± SEM, (n =3), \*\**P <* 0.01, \*\*\*\**P <* 0.0001, Student’s *t* test.

We found Rhes increased the SUMOylation of HDAC1 (Histone Deacetylase 1) and histone 2 B, but it did not affect the SUMOylation of histone H2A.1 (Fig. 3A, B). Surprisingly Rhes decreased the SUMOylation of HNRNPM, PBRM1 and PIASy (PIASy) SUMOylation (Fig. 3B, C). We could not detect SUMOylation of other targets, nucleophosmin (NPM), DGK, and ribosomal protein L26 (RPL26) (Supplementary Fig. 1A). As expected, Rhes was readily SUMOylated under these experimental conditions. Together this data indicates that Rhes differentially modulates the SUMOylation of substrates that are involved in nuclear functions, especially in regulation of gene expression.

### Rhes regulate the expression of genes involved in neuronal morphogenesis in the striatum

As Rhes differentially regulates the SUMOylation of nuclear targets implicated in the gene expression, we hypothesized that Rhes may alter gene expression in the striatum *in vivo*. To investigate this hypothesis, we isolated mRNA from WT and *Rhes*^−/−^mice [RNA from 1 male and 1 female mouse was pooled per sample, triplicate for each group (WT or *Rhes*^−/−^)] and carried out high-throughput RNA seq analysis. We found that Rhes significantly altered the gene expression profile in the striatum (Fig. 4A). Out of ~15,000 genes sequenced, 155 genes are down-regulated, and 52 genes are up-regulated significantly in the striatum of *Rhes*^*−/−*^mice compared to WT control (Fig. 4A, Data file S5). Ingenuity pathway analysis (IPA) showed molecular and cellular functions involved in cell morphology, cellular development, growth and proliferation, and molecular transport are the major hits (Fig. 4B). For example, genes involved in cell morphogenesis; for example, early growth response 2 (Egr2), necdin (Ndn), serum- and glucocorticoid-regulated kinase 1 (Sgk1) and netrin-1 (Ntn1) are upregulated [53–57], and neuropilin1 (Nrp1), basic-helix-loop helix 22 (Bhlhe22) and myocyte enhancer factor 2 (Mef2c) are downregulated [58–62] (Fig 4A). A large majority of the genes, such as ephrin type-A receptor 5 (Epha5) and slit guidance ligand 2 (Slit2), are unaffected. Using qPCR, we further validated selected targets. We confirmed for example, Nrp1, Bhlhe22, Mef2c are downregulated, Egr2 is upregulated, and exostosin glycosyltransferase 1 (Ext1) showed a decreased trend (P**** Value) in the striatum of *Rhes*^*−/−*^mice compare to WT striatum in qPCR analysis (Fig. 4C). This data indicates that Rhes regulates the expression of genes involved in variety of biological process with a prominent role in cellular morphology in the striatum.

**Fig 4.**
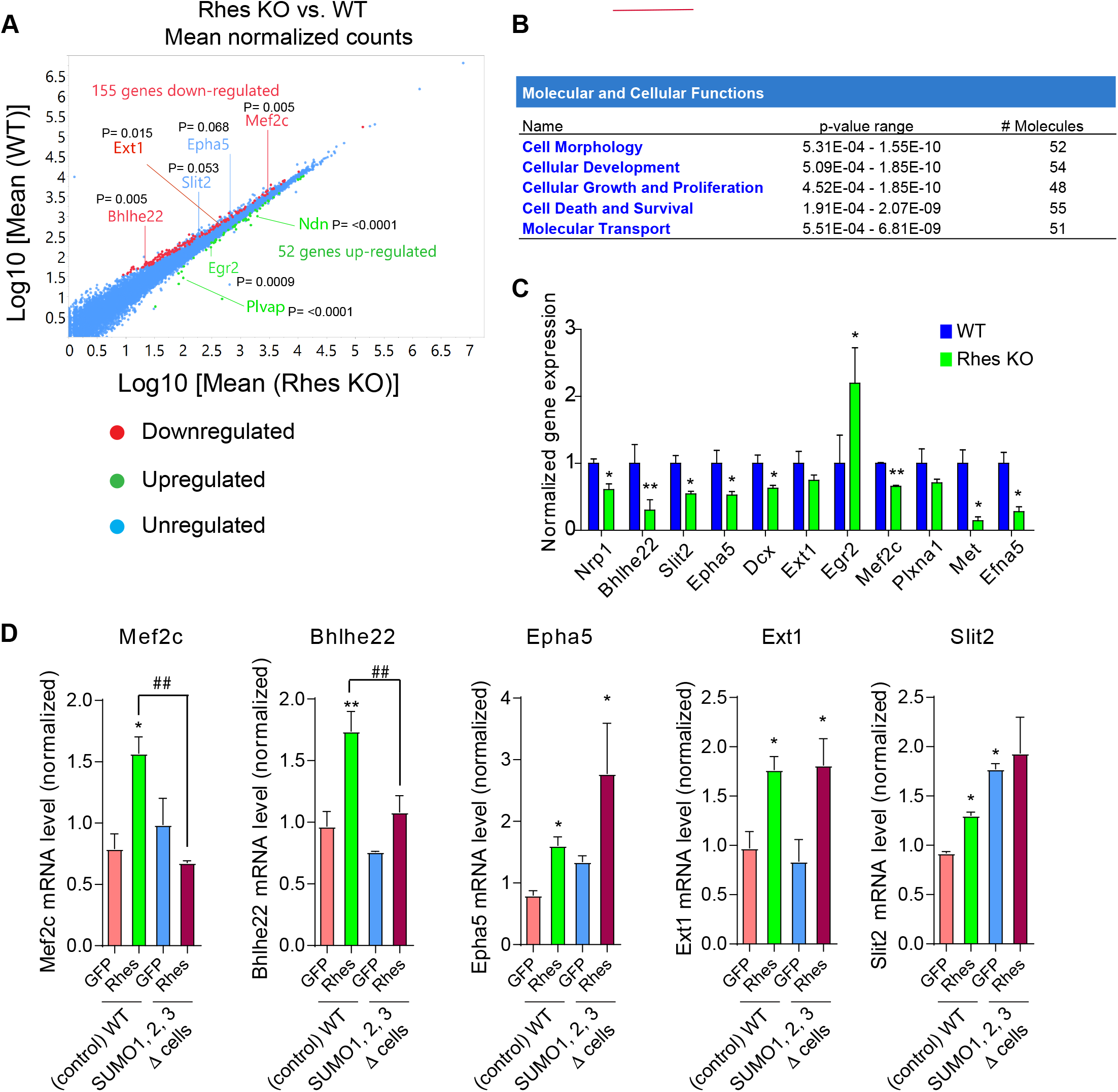
Rhes regulates genes involved in cellular differentiation and morphogenesis via SUMOylation. (**A**) Mean normalized counts of gene expression based on RNA seq data from Rhes KO vs. WT mice striatum. (**B**) IPA analysis of the molecular and cellular functions altered in striatum of Rhes KO mice compared to WT mice from (A). (**C**) Expression of indicated genes (normalized to *Gapdh*), involved in cell morphology and cellular development were validated by qPCR in Rhes KO vs. WT mice striatum. Error bar represents mean ± SEM, (n =3), \**P <* 0.05, **p<0.01 by Student-*t* test. (**D**) Gene expression data for the indicated genes (normalized to *Gapdh*) in WT or SUMO1,2,3 KO (Δ) cells to assess the effect of SUMOylation in presence of GFP or GFP-Rhes. Error bar represents mean ± SEM, (n =3), \**P <* 0.05, \*\**P <* 0.01 compared to GFP in WT. ^##^*P <* 0.01 between Rhes in WT and Rhes in SUMO1,2,3 Δ cells by One-Way ANOVA followed by Newman-Keuls multiple comparison test.

### Rhes regulates a selected gene expression via SUMO

Because Rhes regulates the SUMOylation of proteins such as HDAC1, HNRNPM, PBRM1 and PIASy that are involved in gene expression, [63–68], we investigated if Rhes alters gene expression via SUMO. We employed CRISPR/Cas-9 control (WT cells) and CRISPR/Cas-9 SUMO1/2/3 depleted striatal neuronal cells, that show ~60-70% of loss of SUMO (SUMO Δ cells)[40]. To analyze the effect of Rhes on the expression of Ext1, Mef2c, Slit2, Epha5, and Bhlhe22 in WT and SUMO Δ cells, cells were transfected with GFP or GFP-Rhes and sorted using flow cytometry to obtain enriched population of cells expressing GFP or GFP-Rhes. While Rhes increased the expression of Mef2c, Bhlhe22 in WT cells, it failed to do so in the SUMO Δ cells (Fig. 4D). This result indicates that Rhes positively modulates the gene expression of Bhlhe22 and Mef2c via SUMO. Rhes increased Epha5 and Ext1 in both the control and the SUMO Δ cells, indicating that Rhes promotes Epha5 and Ext1 expression independent of SUMO (Fig. 4D). Furthermore, while Slit2 is induced by Rhes in control cells, a high basal Slit2 expression was found in SUMO Δ cells that was not affected by Rhes expression (Fig. 4D). Collectively, these results indicate Rhes regulates gene expression via SUMO dependent and independent mechanisms.

### Rhes is enriched in the perinuclear membrane fractions

As Rhes mostly regulates gene expression of selected nuclear targets, we investigated if Rhes is localized in the nucleus. Using biochemical tools, we isolated and separated cytoplasmic and nuclear fractions of the striatum from *Rhes*^*+/+*^and *Rhes*^*−/−*^mice (Fig. 5A). Using Western blotting, we confirmed that Rhes is highly enriched in the nuclear fractions that is positive for histone H3 (Fig. 5A). Rhes is also observed in the cytoplasmic fractions that are enriched for cytoplasmic markers, mTOR, mitogen-activated protein kinase kinase (MEK) and lactate dehydrogenase (LDH) (Fig. 5A). Since biochemically isolated subcellular fractions do not unmistakably report the distribution of subcellular proteins, we sought to verify the localization of Rhes using confocal fluorescence microscopy. Imaging analysis revealed that GFP-tagged-Rhes is predominantly associated with perinuclear membranes and it is present at negligible levels in the nucleus (Fig. 5B, red arrows). In addition to perinuclear localization, Rhes is also enriched in the plasma membrane (Fig. 5B, yellow arrowhead), and membranous vesicles, as consistent with our previous reports [40, 69]. Altogether, our result indicates that Rhes is predominantly localized on the perinuclear membranes. We propose that Rhes may affect SUMOylation of nuclear targets in the perinuclear location (Fig. 5C).

**Fig 5.**
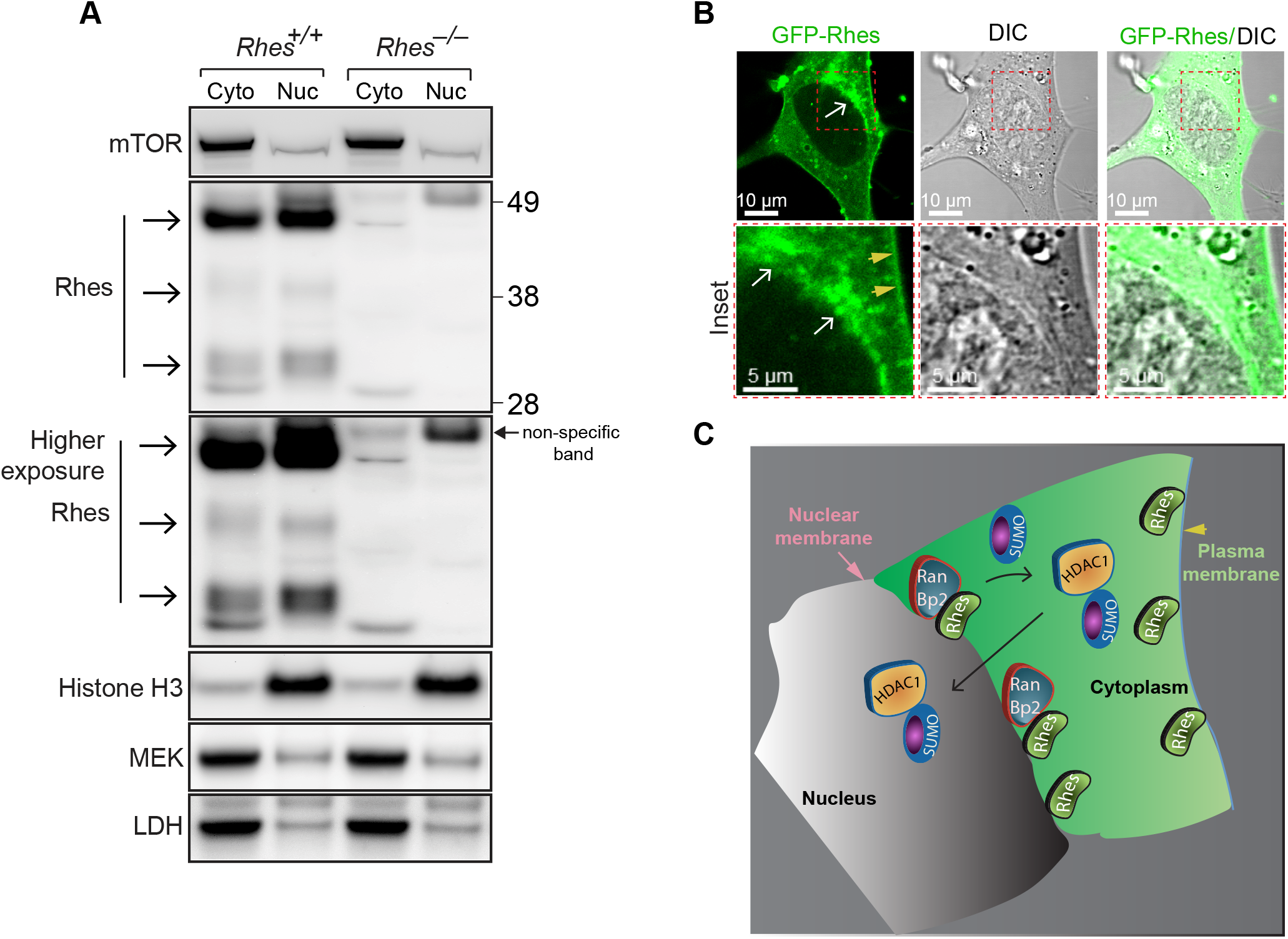
Rhes is preferentially localized around the perinuclear membrane. (**A**) Subcellular localization of Rhes was assessed in cytosolic and nuclear fractions from Rhes KO (*Rhes^−/−^*) vs. WT (*Rhes^+/+^*) mice striatum. The cytosolic markers MEK and LDH and the nuclear marker Histone H3 were probed to validate proper cellular fractionation. (**B**) Representative confocal and brightfield (DIC) image of striatal neuronal cell expressing GFP-Rhes, indicating the localization of Rhes on nuclear membrane and perinuclear space. Inset shows the boxed area at higher magnification. White arrows indicate perinuclear localization of Rhes, and yellow arrowhead indicates the localization of Rhes on plasma membrane. (**C**) Model depicting that Rhes can SUMOylate its targets, including HDAC1, in the cytoplasm and promotes its entry into the nucleus to regulate gene expression.

## DISCUSSION

The data presented here demonstrate that a) Rhes is SUMOylated on multiple lysine sites; b) Rhes regulates the SUMOylation of nuclear proteins, and c) Rhes regulates gene expression in the striatum at least partly via SUMO-mediated mechanisms. Importantly, despite that, it is well known that Rhes is post-translationally modified *in vivo* [9, 70]; however, the nature of modification remained unknown. This study indicates that SUMO may contribute to such *in vivo* Rhes modification.

Interestingly, the presence of multiple closely spaced SUMO modification sites on Rhes on K110, K114, K120, K124 indicates that these residues may act as anchors that can associate with SUMO-interacting motif (SIM) which are known to be involved in protein-protein interaction [71, 72]. Such closely spaced SUMOylation events are also found on other SUMO E3 ligases such as PIAS1 lysines K40, K46, K56, and K58 and PIASy lysines K59 and K69, K128 and K135; and RanBP2 K1596 and K1605, K2513 and K2531, and K2571 and K2592 [73, 74]. Thus, multiple lysine SUMO modifications appear to be a characteristic feature across many SUMO E-3 like proteins. Identifying the role of each lysine modification, and whether Rhes auto-SUMOylates itself or if other potential SUMO E3 ligases SUMOylate Rhes needs further investigation.

Intriguingly, we report that Rhes differentially alters the SUMOylation of SUMO E3 ligases. Rhes decreased the overall SUMOylation on PIAS in the cells (Fig. 3B), while promoting the specific SUMOylation of PIASy on lysine 59 (Fig. 1H). Similarly, MS analysis revealed that Rhes significantly increases the SUMOylation of RanBP2 on lysine 1605, but not on lysine 2592 (Fig 2A, Data file S5). These results indicate that Rhes may differentially affect the SUMOylation of SUMO E3 ligase(s) on selected lysine targets. In the future, it will be interesting to test whether PIASy or RanBP2 can act as SUMO E3 ligase for Rhes. Even though it is known that SUMOylation modifies SUMO E3 ligases, the biological significance of such modifications remains less understood. Previously, we reported that Rhes promoted cross-SUMOylation between SUMO E1 (Aos/Uba9) and E2 (Ubc9) ligases and predicted such regulation might work as “symbiotic” regulation between two evolutionarily conserved SUMO E1 and E2 enzymes [44]. Based on the data presented here, we propose that Rhes may also promote reciprocal and symbiotic regulations between SUMO E3 ligases through SUMO modification that may have a significant impact on regulating complex cellular and behavioral functions of the striatum via protein-protein interactions [9].

The results presented here clearly indicate that Rhes is involved in the regulation of gene expression in the striatum, intriguingly, it can both increase and decrease gene expression in the striatum. A previous independent study also reported that Rhes might inhibit gene expression by acting as a cis-modulator [75], but the mechanisms were unclear. It is well established that SUMOylation participates in cis-regulation and is involved in both transcriptional repression as well as activation functions [51, 76, 77]. As Rhes KO cells show overall diminished SUMOylation in the striatum [44] and altered gene expressions, we predict that the Rhes-SUMO pathway may regulate gene expression as cis-modulator, for example via the SUMOylation of transcription factors. Consistent with this notion, although we did not observe a significance alteration of HDAC1 SUMOylation on lysine 89 or lysine 476 in our proteomics analysis (Data S3), presumably due to low stoichiometry, we found that Rhes enhances the SUMOylation of HDAC1 in our biochemical experiments (Fig. 2B). Thus Rhes may alter HDAC1 activity via SUMOylation at lysine 89 and lysine 476, a catalytically essential residue involved in the gene repression [78–80].

Previous studies indicate that SUMOylation of H2B is involved in the repression of gene expression [81]. Because we found that Rhes increases the SUMOylation of H2B (Fig. 3), which is SUMOylated at multiple lysine (Data file S4), including previously reported, lysine 6 [81], we propose that Rhes-SUMO pathway may affect gene expression via more than one nuclear targets.

The cellular compartment in which Rhes regulates the SUMOylation of nuclear proteins remains to be elucidated. Since we found that Rhes is enriched on the perinuclear space (Fig. 5B), we predict that this location may serve as the prime site for SUMOylation of proteins (Fig. 5C). In support of this notion, previous studies showed that SUMO E3 ligase RanBP2 localized on the perinuclear location [the cytoplasmic side of nuclear pore complex (NPC)], which regulates import and export of proteins, and mediates global gene expression in cell models [82, 83]. RanBP2 also enhances the SUMOylation of HDAC4 and HNRNPM proteins, which are involved in mRNA splicing and transport [84, 85]. Although the localization of Rhes on the NPC is not currently documented, we found that Rhes coimmunoprecipitates with RanBP2 and Sec13, two known components of the NPC. Rhes has been shown to interact with NPC associated proteins during motor stimulation in the striatum, including HNRNP (L1 and H2 isoforms) [9]. Thus, we predict that Rhes may associate with the NPC to regulate the SUMOylation of targets involved in gene regulation as well as mRNA splicing, a process shown to require SUMOylation [86]. In addition, Rhes may alter SUMO-independent signaling, such as the modulation of mTOR and PKA signaling pathways that are linked to gene expression [6, 8, 87–89].

Our finding demonstrates that Rhes alters the expression of genes involved in cell morphology and differentiation (Fig. 4) and is a potent inducer of TNT-like cellular protrusions [40]. We propose that Rhes may mediate TNT-like protrusions and cell-to-cell transport of cargoes via the regulation of the expression of transcription factors such as Mecf2, Egr2, Bhlhe22, which are known regulators of neuronal morphology and differentiation in a SUMO dependent manner. Consistent with this notion, depletion of SUMO diminishes the formation of TNT-like protrusions [40] and cell-to-cell transport as well as alters Rhes mediates expression of Mef2c and Bhlhe22-1 9 (Fig. 4).

Collectively, by combining high throughput interactomics, SUMO proteomics, gene expression analysis, and biochemical tools, we demonstrate that Rhes promotes the SUMOylation of nuclear substrates that are involved in gene expression. Thus, Rhes may impact striatal function and dysfunction associated with neurological and neurodegenerative disease via the SUMO-mediated regulation of gene expression.

## MATERIALS AND METHODS

### Reagents, Plasmids and Antibodies

Unless otherwise noted, reagents were obtained from Sigma. Full length Rhes cDNA constructs, Myc, GST or GFP tagged, were cloned in pCMV vectors (Clontech) [11, 12, 40].. Mass spectrometry detection compatible His6-mSUMO1, His6-mSUMO2 and His6-mSUMO3 were cloned as described [90, 91]. p181 pK7-HDAC1 (GFP) (Addgene; 11054) were from Ramesh Shivdasani. H2B-mCherry (Addgene; 20972) were from Robert Benezra. mH2A1.2-CT-GFP (Addgene, 45169) were from Brian Chadwick, Hunt Willard. pT7-V5-SBP-C1-HshnRNPM (Addgene; 64924) were from Elisa Izaurralde. GFP-PBRM1 (Addgene; 65387) were from Kyle Miller. Flag-hPIASy (Addgene; 15208) were from Ke Shuai. GFP-NPM WT (Addgene; 17578) were from Xin Wang. DGK beta (Addgene; 35405) were from Robert Lefkowitz and Stephen Prescott. pCMV-L26-Flag (Addgene; 19972) were from Moshe Oren. 6x-His Tag Antibody (clone HIS.H8) (1:1000, # MA1-21315) was from ThermoFisher Scientific. Antibodies for GST-HRP (1:5000, #sc-138), and Myc (1:3000, #sc-40) were obtained from Santa Cruz Biotechnology. mCherry antibody (1:1000, NBP2-25157) was from Novus Biologicals. Flag antibody (1:1000, F7425) was obtained from Sigma-Aldrich. HA.11 Epitope Tag Antibody (1:1000, 901513) was from BioLegend (previously Covance catalog# MMS-101R). V5-Tag (1:1000, #13202), GFP (1:1000, #2956), mTOR (1:3000, #2983), Histone H3 (1:10,000, #4499), MEK1/2 (1:1000, #8727) and LDH (1:5000, #2012;) were from Cell Signaling Technology, Inc. Rhes antibody (1:1000, RHES-101AP) was from Fabgennix.

### Ni-NTA Denaturing Pull Down

Ni-NTA pull down of His-SUMO conjugates was performed as previously described [12, 92]. Briefly, HEK293 cells (expressing transfected His-SUMO mutant and indicated constructs) were pretreated with MG132 (25 μM, 4 hr), rinsed in PBS, scraped from 10cm^2^ dishes, and centrifuged at 750 x g for 5 min. Cell pellets were then directly lysed in Pull Down Buffer [6 M Guanidine hydrochloride, 10 mM Tris, 100 mM sodium phosphate, 40 mM imidazole, 5 mM β-mercaptoethanol (β-ME), pH 8.0] and sonicated. Lysates were then clarified by centrifugation at 3,000 × g for 15 minutes. All subsequent wash steps were performed with 10 resin volumes of buffer followed by centrifugation at 800 x g for 2 min. Ni-NTA Agarose beads (#30210; Qiagen) were pre-equilibrated by washing three times with Pull Down Buffer. After equilibration, beads were resuspended in Pull Down Buffer as a 50% slurry of beads to buffer. After protein quantification of cell lysates, 1 mg of lysate was added to 40 μL of Ni-NTA bead slurry to a total volume of 1 mL in microcentrifuge tubes. Beads were then incubated overnight at 4°C mixing end over end. The following day, beads were centrifuged and underwent washing once in Pull Down Buffer, once in pH 8.0 Urea Buffer (8 M Urea, 10 mM Tris, 100 mM sodium phosphate, 0.1% Triton X-100, 20 mM imidazole, 5 mM β-ME, pH 8.0), and three additional times in pH 6.3 Urea Buffer (8 M Urea, 10 mM Tris, 100 mM sodium phosphate, 0.1% Triton-X100, 20 mM imidazole, 5 mM β-ME, pH 6.3). Elution was performed using 20 μL of Elution Buffer (pH 8.0 Urea Buffer containing 200 mM imidazole, 4X NuPAGE LDS loading dye, 720 mM β-ME). Samples were then heated at 100°C for 5 minutes and directly used for Western Blotting. Inputs were loaded as 1% of the total cell lysate. The SUMOylation is quantified by normalizing the intensity of SUMOylation bands to the respective unmodified bands using Image-J.

### GFP-Rhes localization studies

Approximately 75,000 STHdh^Q111/Q111^ cells were seeded on 35mm glass bottom dishes. After 24h the cells were transfected with indicated plasmids. Cells were imaged live using a Zeiss 880 confocal microscope at 63X oil immersion Plan-apochromat objective (1.4 NA).

### Purification of SUMOylated Rhes interacting proteins

Proteins extracts from GST + mSUMO3, and GST-Rhes + mSUMO3 expressing HEK298 cells were purified with glutathione beads to recover global Rhes interacting partners. The resulting purified material was subjected to denaturing Ni-NTA purification step to enrich for the SUMOylated interacting parteners. The Ni-NTA bound material was digested with trypsin, desalted on C-18 and analyzed by LC-MS, as described before [47, 93, 94].

### SUMO peptide enrichment and MS detection

SUMOylated proteins were enriched from myc + mSUMO3 or myc-Rhes + mSUMO3 expressing HEK293 cells using Ni-NTA. The solid support bound material was digested on the Ni-NTA beads with trypsin and the resulting peptides desalted on C-18 cartridges. SUMOylated peptides were immunopurified with the anti-NQTGG antibody that recognizes the NQTGG remnant exposed on the SUMOylated lysine upon tryptic digestion. SUMO site quantification and quantification was carried out with MaxQuant, as described before [46, 94, 95].

#### SUMO1/2/3 Knockout in striatal Cells

Striatal cells deleted for SUMO1, 2 and 3 is using CRISPR/Cas9 SUMO gRNAs as described [40].

#### Mice

For fractionation experiments, we used Rhes KO (*Rhes*^−/−^) mice, and C57BL/6J mice. Rhes KO mice were obtained from Alessandro Usiello and were backcrossed with C57BL/6J mice at least 8 generations; homozygous Rhes KO were used for all of the experiments [9, 13]. WT mice (C57BL/6) were obtained from Jackson Laboratory and maintained in our animal facility according to Institutional Animal Care and Use Committee at The Scripps Research Institute.

### Nuclear and Cytoplasmic Separation from striatum

Striatum from C57BL/6J and Rhes KO mice was fractionated using the nuclear/cytosol fractionation kit according to the manufacturer’s instructions with minor modifications (BioVision). Briefly, striatum from C57BL/6J and Rhes KO mice was rapidly dissected out and homogenized in CEB (cytosolic extraction buffer)-A with DTT and protease inhibitor, and incubating for 10 min on ice prior to addition of CEB-B. The lysates were vortexed for 5 sec and incubated on ice for 1 min. The lysates were then centrifuged at 4°C for 5 min at 16,000 × g in a microcentrifuge. And the supernatants were kept as the cytoplasmic fraction. The nuclear pellet was resuspended in NEB (nuclear extraction buffer). And vortexed the lysates for 15 sec. This step was repeated 5 times every 10 min. The nuclear pellet was centrifuged at 4°C for 10 min at 16,000 × g in a microcentrifuge. And the supernatants were kept as the nuclear fraction. The protein concentration was determined in the cytoplasmic and nuclear fractions using the BCA Protein Assay Kit (Pierce, Rockford, IL, USA). Equivalent amounts of protein samples (50 μg) were resolved by SDS-PAGE followed by immunoblotting as described below.

### Western blotting

Equal amounts of protein (20-50 μg) were loaded and were separated by electrophoresis in 4 to 12% Bis-Tris Gel (Thermo Fisher Scientific), transferred to polyvinylidene difluoride membranes, and probed with the indicated antibodies. HRP-conjugated secondary antibodies (Jackson ImmunoResearch Inc.) were probed to detect bound primary IgG with a chemiluminescence imager (Alpha Innotech) using enhanced chemiluminescence from WesternBright Quantum (Advansta). Where indicated the membranes were stained for ponceau.

### Global mRNA-seq from WT and Rhes KO striatum

WTor Rhes KO mice striatum [1 male and 1 female (pooled), in triplicate were lysed in Trizol reagent. RNA was extracted from miRNeasy kit from Qiagen (cat. No. 217004) and mRNA-seq was performed as described before [96].

### qPCR validation of targets

Striatum of mice (WT or Rhes KO) lysed in Trizol reagent. 250 ng RNA was used to prepare cDNA using Takara primescript kit (Cat no. 6110A) using random hexamers. The qRT-PCR of genes was performed with SYBR green (Takara RR420A) reagents. Primers for all the genes were designed based on sequences available from the Harvard qPCR primer bank. The primer sequences are as follows:

*Gapdh* mouse (Forward primer) 5’ primer AGGTCGGTGTGAACGGATTTG (Reverse primer) 3’ primer TGTAGACCATGTAGTTGAGGTCA
*Nrp1* mouse (Forward primer) 5’ primer GACAAATGTGGCGGGACCATA (Reverse primer) 3’ primer TGGATTAGCCATTCACACTTCTC
*Bhlhe22* mouse (Forward primer) 5’ primer TGAACGACGCTCTGGATGAG (Reverse primer) 3’ primer GGTTGAGGTAGGCGACTAAGC
*Slit2* mouse (Forward primer) 5’ primer GGCAGACACTGTCCCTATCG (Reverse primer) 3’ primer GTGTTGCGGGGGATATTCCT
*Epha5* mouse (Forward primer) 5’ primer AAGGAACCCTGTGGCTATTGG (Reverse primer) 3’ primer GCAAACATGCCCGTTTTAGAGAA
*Dcx* mouse (Forward primer) 5’ primer CATTTTGACGAACGAGACAAAGC (Reverse primer) 3’ primer TGGAAGTCCATTCATCCGTGA
*Ext1* mouse (Forward primer) 5’ primer TGGAGGCGTGCAGTTTAGG (Reverse primer) 3’ primer GAAGCGGGGCCAGAAATGA
*Egr2* mouse (Forward primer) 5’ primer GCCAAGGCCGTAGACAAAATC (Reverse primer) 3’ primer CCACTCCGTTCATCTGGTCA
*Mef2c* mouse (Forward primer) 5’ primer ATCCCGATGCAGACGATTCAG (Reverse primer) 3’ primer AACAGCACACAATCTTTGCCT
*Plxna1* mouse (Forward primer) 5’ primer GGGTGTGTGGATAGCCATCAG (Reverse primer) 3’ primer GCCAACATATACCTCTCCTGTCT
*Met* mouse (Forward primer) 5’ primer GTGAACATGAAGTATCAGCTCCC (Reverse primer) 3’ primer TGTAGTTTGTGGCTCCGAGAT
*Efna5* mouse (Forward primer) 5’ primer ACACGTCCAAAGGGTTCAAGA (Reverse primer) 3’ primer GTACGGTGTCATTTGTTGGTCT

### Bioinformatics and statistical analysis of data analysis of proteomics

Normalized gene counts were averaged and log10 transformed for WT and Rhes KO samples and were plotted against each other where Rhes Ko log10 mean values were on x-axis and WT log10 mean values were on y-axis. Differentially regulated genes were identified using padj <0.05 cut off and up (higher in Rhes KO) and down (higher in WT) regulated genes were marked with green and red respectively. The graph was generated using JMP®, Version 13.2.1. SAS Institute Inc., Cary, NC.

### Statistical Analysis

Data were expressed as means ± SEM. All of the experiments were performed at least in triplicate and repeated twice at minimum. Statistical analysis was performed using Student’s *t* test or one-Way ANOVA followed by Newman-Keuls multiple comparison test (GraphPad Prism 7).

### Data availability

Sequencing data have been submitted to the Gene Expression Omnibus (GEO) data repository, under the accession number GSE150990.

## LEGENDS

**Supplementary Figure 1.**
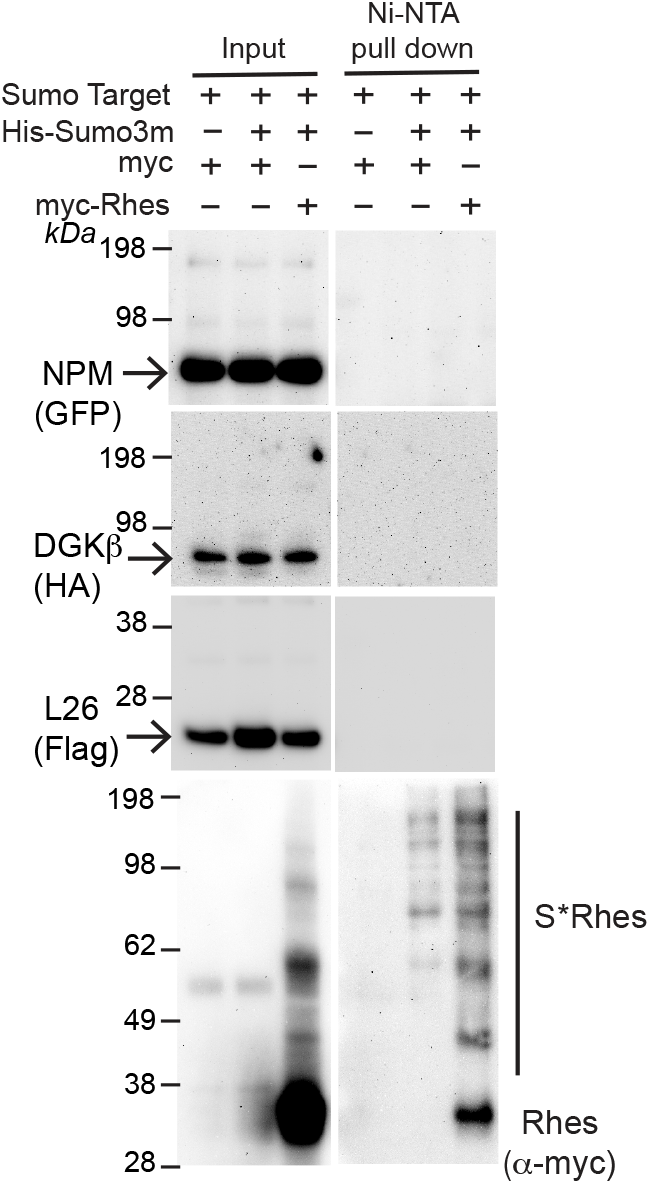
Western blot of Ni-NTA enrichment of SUMOylated proteins from lysates obtained from HEK293 cells expressing myc + His-mSUMO3 or myc-Rhes + His- mSUMO3 constructs either with GFP-NPM, HA-DGK-β or Flag-L26.

## Author contributions

S. Su conceptualized and designed the project. O.R carried out all the biochemical SUMO work. M. S carried out mRNA seq, qPCR, SUMO-KO cell generation and confocal microscopy. N.S carried out biochemical separation of Rhes. UNRJ assisted in mRNA seq and cell culture experiments. F. M and T.P designed the mass spec (MS) compatible his-mSUMO3 construct and carried out MS analysis of Rhes interactors and identification of SUMO. G.C assisted in bioinformatics. P.K generated mRNA seq library and sequencing. S.Su wrote the paper with input from the co-authors.

## Acknowledgement

We would like to thank Melissa Benilous for her administrative help and the members of the lab for their continuous support and collaborative atmosphere. This research was supported by funding from National Institutes of Health/National Institute of Neurological Disorders and Stroke grant R01-NS087019-01A1, National Institutes of Health/National Institute of Neurological Disorders and Stroke grant R01-NS094577-01A1, and a grant from Cure Huntington Disease Initiative (CHDI) foundation.

## Notes

### Competing Interest Statement

The authors have declared no competing interest.

